# Efficient Sex Separation by Exploiting Differential Alternative Splicing of a Dominant Marker in *Aedes aegypti*

**DOI:** 10.1101/2023.06.16.545348

**Authors:** Shih-Che Weng, Igor Antoshechkin, Eric Marois, Omar S. Akbari

## Abstract

Only female mosquitoes consume blood and transmit deadly human pathogens. Therefore, it is critical to remove females before conducting releases for genetic biocontrol interventions. Here we describe a robust sex-sorting approach termed SEPARATOR (<u>S</u>exing <u>E</u>lement <u>P</u>roduced by <u>A</u>lternative <u>R</u>NA-splicing of <u>A</u> <u>T</u>ransgenic <u>O</u>bservable <u>R</u>eporter) that exploits sex-specific alternative splicing of an innocuous reporter to ensure exclusive dominant male-specific expression. Using SEPARATOR, we demonstrate reliable sex selection from larval and pupal stages in *Aedes aegypti*, and use a Complex Object Parametric Analyzer and Sorter (COPAS®) to demonstrate scalable high-throughput sex-selection of first instar larvae. Additionally, we use this approach to sequence the transcriptomes of early larval males and females and find several genes that are sex-specifically expressed in males. SEPARATOR can simplify mass production of males for release programs and is designed to be cross-species portable and should be instrumental for genetic biocontrol interventions.

## Introduction

The mosquito is the most deadly animal on the planet, estimated to have killed roughly 50% of humans since the dawn of humanity^1^. This year, roughly 700 million cases of mosquito transmitted diseases worldwide shall occur resulting in over a million deaths^2^. Insecticides and larvicides have traditionally been used to control mosquitoes, however mosquitoes have evolved resistance^3–7^, and harmful chemicals are unfortunately poisoning our environment and killing non-target species^8–10^. Moreover, a warming global climate is expanding habitable ranges of mosquitoes and their pathogens^11–13^. Consequently, the number of people at risk of contracting mosquito transmitted pathogens continues to rise, indicating innovative technologies are critical to combat this global threat.

To genetically suppress mosquito populations, several biocontrol strategies have been developed including the classical sterile insect technique (SIT)^14^, release of insects carrying a dominant lethal (RIDL)^15^, female specific RIDL (fsRIDL)^16–20^, Wolbachia-based incompatible insect technique (IIT)^21^, the precision guided sterile insect technique (pgSIT)^22^, Ifegenia (inherited female elimination by genetically encoded nucleases to interrupt alleles)^23^, and homing-based suppression gene drives^24^. The ultimate goal for each of these approaches is to scalably release modified mosquitoes into the environment to safely achieve sustained species-specific population suppression. Given that only female mosquitoes consume blood and transmit pathogens, exclusive release of male-only cohorts is important. While some of these approaches have built-in self-sexing mechanisms (e.g. gene drives^24^, fsRIDL^20^, Ifegenia^23^, pgSIT^22^), others require laborious manual separation of females prior to release (e.g. classical SIT, Wolbachia-based IIT^21^) significantly reducing scalability. Moreover, for Wolbachia-based IIT any accidental female releases can undermine the intervention ^25, 26^, underscoring the imminent need for efficient and reliable sexing.

Sex-sorting mosquitoes can be done manually by hand, however this is laborious, error-prone, and not scalable for genetic biocontrol interventions. In *Aedes* and *Culex* mosquitoes, sexual dimorphism of pupa size can aid sex separation ^27, 28^, however pupal size is density-dependent requiring optimized rearing conditions. To overcome this limitation, AI-assisted optical adult sex-sorting systems have been developed that can differentiate adults based upon sexual dimorphic features with impressive accuracy^29^. However, these have only been developed in *Aedes aegypti* and require sorting at the fragile adult stage which is not ideal, slow, and less scalable. Alternatively, selectable markers have been genetically linked to sex chromosomes in *Anopheles*^30–34^, or to sex-determining loci in *Aedes*^35–37^. However, genetic linkage is often broken by meiotic recombination, or translocations, making these strains less stable when scaled ^14, 38^. Strains have also been developed using sex-specifically expressed promoters in mosquitoes ^19, 39–41^ to express fluorescent proteins in the gonads, however given the small size of the gonads the expression is often weak and stage-specific^42^. There have also been efforts to exploit sex-specific alternative splicing (SSAS) to enable red fluorescent protein (RFP) expression in the flight muscles in *Aedes*^17^, or to bias green fluorescent protein (GFP) expression in larvae in *Anopheles*^43, 44^. However, these sexing strains are engineered using species-specific components making them less portable across species. Additionally, the expression patterns are often not strong, or early enough, to allow for reliable sorting in initial larval stages which would be ideal for genetic biocontrol interventions. Taken together, the development of simple, stable, and reliable sex-sorting technologies that can be facilely engineered in multiple mosquito species and enable early larval sex sorting remains to be developed.

Here we engineer an innovative sex-sorting approach that: (i) exploits male specific expression via sex-specific alternative splicing (SSAS) of an innocuous bright fluorescent marker; (ii) enables sex-sorting during early larval development and beyond; (iii) is adaptable for high-throughput sorting; (iv) does not rely on sex-chromosome linkage; (v) is genetically stable and not prone to breakage by meiotic recombination or chromosomal rearrangements; (vi) enables positive selection of males to further increase robustness; and (vii) is facilely portable to alternate species as it utilizes transposable elements and promoters and markers that are cross-species portable. We call our approach SEPARATOR (<u>S</u>exing <u>E</u>lement <u>P</u>roduced by <u>A</u>lternative <u>R</u>NA-splicing of <u>A</u> <u>T</u>ransgenic <u>O</u>bservable <u>R</u>eporter). SEPARATOR exploits SSAS of a widely expressed fluorescent reporter to provide positive selection of males that can be integrated into genetic biocontrol interventions. Using SEPARATOR, we demonstrate positive identification of male larvae in *Aedes* mosquitoes, as early as mature embryos and beyond. By harnessing the capabilities of COPAS, we demonstrate sorting of over 100,000 GFP+ males with speeds of up to 740 larvae/minute, achieving a GFP-negative female contamination rate between 0.01% to 0.03%. Taken together, this method provides a much needed tool to scale and reliably release males for genetic biocontrol interventions.

## Results

### Engineering SEPARATOR

To generate SEPARATOR a sex-specific alternatively spliced intron derived from the *Ae. aegypti doublesex* (*AaeDsx*) gene was utilized (**Fig. 1A and fig. S1A**). Dsx is a highly conserved transcription factor involved in sex determination of insects. In *Ae. aegypti* the male specific *AaeDsx* intron is ∼26.5 kb which is a bit large to work with. Therefore, we truncated this intron by preserving splicing factor binding sites including Tra/Tra-2 and RNA binding protein 1 (RBP1) binding sites to retain the sex-specificity of this intron ^45^. This resulted in a smaller *AaeDsx* intron of 4.5 kb in size (**Fig. 1A and fig. S1A**). The reading frame was initiated by adding a start codon with a Kozak sequence, expressed using a constitutive *Hr5IE1 Ac*MNPV baculovirus promoter previously shown to work in many species ^46–53^. To open the reading frame, nine stop codons located in endogenous exon 5b were excluded (**table S6**). The coding sequences for EGFP and DsRed were strategically designed to overlap, allowing for their expression in a sex-specific manner. The DsRed coding sequence was designed to be in-frame with a female-specific product (exon4, engineered exon5b, and exon6) to control female-specific DsRed expression. In addition, the male-specific splicing product, involving exon 4 and exon 6, was designed to be in-frame with the EGFP coding sequence (**fig. S1**).

**Figure 1.**
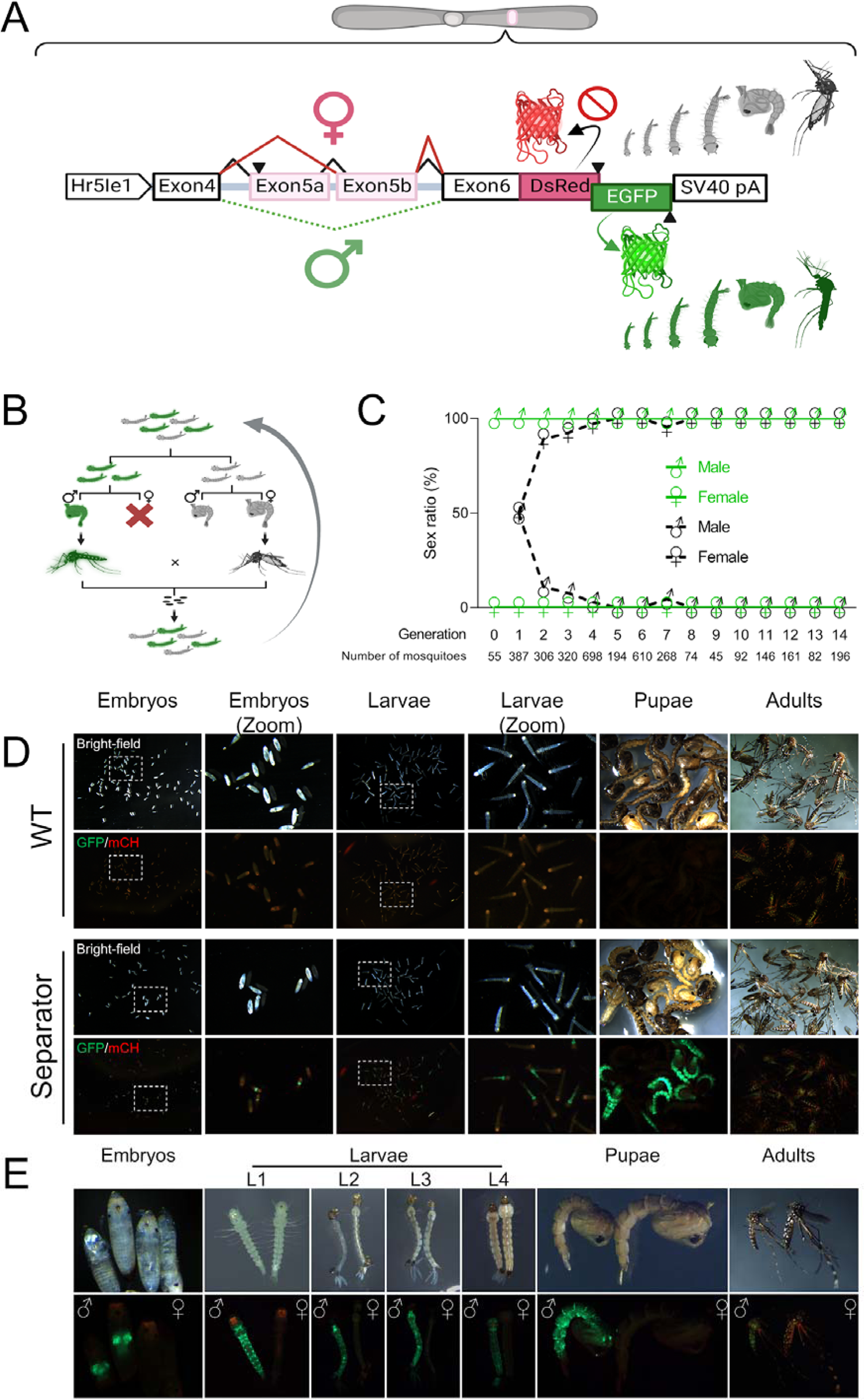
SEPARATOR in *Ae. Aegypti*. (A) The sex-specific splicing module of AaDsx was used to construct SEPARATOR in *Ae. aegypti.* The expression of SEPARATOR was driven by the constitutive baculovirus promoter, Hr5Ie1. The male-specific splicing product was in-frame with the EGFP coding sequence, while inclusion of stop codons in exon 5b prevented in-frame expression of the DsRed in females. The SV40 pA served as the polyadenylation signal. The construct is not to scale. (B) Strategy for generating homozygotes. GFP-positive larvae were sorted and the sex of each was determined at the pupal stage. GFP-positive males were then crossed with GFP-negative females in order to produce homozygotes. (C) GFP-positive mosquitoes are male exclusively. Mosquitoes were sorted by their GFP signal at the larval stage and used a microscope to examine the sex ratio based on the morphological differences in genital lobe shape at pupal stage, which are specific to each sex. (D) The embryo, larva, pupa, and adult stages of wild-type (Liverpool) and SEPARATOR mosquitoes were collected and photographed using a fluorescent stereomicroscope (Leica M165FC). Eggs, aged between 24-48 hours after being laid, were subjected to a 15-30 minute treatment with 30% NaOCl solution (containing approximately 3.6% active chlorine at the final concentration) to remove the chorion and enable visualization of the embryo. Eggs were hatched in deionized water within a vacuum chamber, and the resulting hatched larvae were then collected as L1 larvae. (E) The developmental stages of mosquitoes were collected and photographed using a fluorescent stereomicroscope (Leica M165FC). The images consist of two panels: the upper panel displays the bright-field images, while the lower panel showcases the GFP/mCH channel images.

The SEPARATOR construct was then introduced into the mosquito genome to generate a genetic sex-sorting strain via the piggyBac transposon. The intended plan was to ensure that all mosquitoes expressing GFP would be male, while those expressing DsRed would be female. Interestingly, the results following microinjection revealed that all 55 EGFP-expressed larvae were male at the pupal stage in G0 (**table S1**). However, no DsRed-expressed larvae were observed in G0. All the pupae from G0 were sexed and resulting adults were crossed with wild-type mosquitoes, and stable transgenic lines were selected by the fluorescence marker (G1). In G1-onward, similar results were observed, where 100% of the EGFP-positive larvae were male.

To validate the sex-specific splicing pattern of the *AaeDsx* splicing module, a comprehensive analysis was carried out utilizing reverse transcription polymerase chain reaction (RT-PCR) and RNA sequencing. A total of fifty EGFP-positive L1 larvae and fifty EGFP-negative L1 larvae were collected for further analysis. To perform RT-PCR, we utilized primers designed to target the 3’ end of the *Hr5IE1* promoter and the 5’ end of the EGFP sequence (**fig. S1A**). Subsequently, Sanger sequencing was carried out to analyze the obtained PCR products. Our results showed that the utilization of truncated intron 4, engineered exon 5b, intron 6, and exon 6 sequences in RNA splicing exhibited sex-specificity (**fig. S1B**). Both Sanger sequencing and RNA sequencing (RNA-seq) analysis yielded comparable results, aligning with our anticipated splicing patterns (**fig. S1B and fig. S2**). The sex-specific RNA splicing pattern indicated the female splicing product being in-frame with the DsRed coding sequence, while the male splicing product resulted in (−1) frameshift, leading to the DsRed coding sequence being out of the frame and in-frame with the EGFP coding sequence. Interestingly, both the RT-PCR and RNA-seq analyses revealed that the predominant products observed in females comprised exon4, exon5b, and exon6 (**fig. S1B and C and table S3**). Notably, these exons were found to be in-frame with the DsRed coding sequence (**fig. S1B and fig. S2**). Therefore, it can be inferred that the expression level of transcripts specific to females and the splicing pattern of female-specific products do not present any obstacles to DsRed expression. However, DsRed signals were not observed in the transgenic larvae.

We determined transgene integration sites (**fig. S3**) and generated a homozygous line. To do this, larvae were sorted by fluorescence and separated into two groups, EGFP-positive and EGFP-negative, then the sex ratio of these two groups was examined at the pupal/adult stage. The EGFP-positive males were crossed to EGFP-negative females to enrich for homozygotes (**Fig. 1B**). In 15 generations, we manually screened a total of 3635 EGFP+ larvae and confirmed the resulting sex at the pupa and adult stage. Remarkably, 100% of the GFP+ larvae were male (**Fig. 1C and table S1**). In summary, the results demonstrate that the SEPARATOR technology is an efficient and effective way of separating male (EGFP+) and female mosquitoes (EGFP-), which could have important implications for broad adaptation of SIT for mosquito control.

### Measuring Efficacy of Sex Sorting at Scale

To evaluate the applicability of SEPARATOR during the entire life cycle of mosquitoes, we proceeded to explore the timing of EGFP expression. We found that strong EGFP signals were expressed from late embryo all the way through to the adult stage of the mosquitoes (**Fig. 1D and E**). In addition, when compared to the wild-type control (Liverpool), the EGFP intensity of SEPARATOR mosquitoes is strong enough to differentiate between EGFP-positive (male) and EGFP-negative (female) in a pooled condition from the early stage of mosquito’s life cycle (**Fig. 1D**). Based on these results, it can be concluded that SEPARATOR is a powerful and robust system for mosquito sex sorting.

To evaluate the suitability of SEPARATOR for automated sex sorting, we conducted fluorescence-based flow cytometry sorting on batches of several thousand first instar larvae using COPAS. This generated a fluorescence diagram where male larvae expressing EGFP formed a distinct cluster, clearly separate from the GFP negative female larvae (**Fig. 2 and fig. S4**). The EGFP-positive larvae were selected and sorted in pure mode at flow rates ranging from 20 to 70 objects per second. Even at this high flow rate, we were able to recover 70-80% of the EGFP-positive larvae, resulting in sorting speeds of 740 larvae/minute **table S2**). In total, 108,570 larvae were sorted using COPAS from generations G7 to G9. After a single sorting, a single instance of 0.1 to 0.45% contamination of EGFP-negative (female) larvae was observed within the male population (**table S2**). While slower flow rates could potentially yield a more complete recovery of males and minimize or eliminate contamination with females, the high sorting speed represents a compromise between recovery rates and the production speed required for mass production^54^. To address the issue of female contamination, additional measures were implemented. Through quality control sorts, the contamination rate of EGFP-negative larvae (females) was successfully reduced to 0.01-0.03% (**table S2**). This was accomplished by subjecting the sorted larvae, a total of 78,861 individuals at G9, to a second round of sorting using the COPAS. This second sorting phase specifically targeted objects that deviated from the EGFP-positive gate, employing the “enrich mode” for enhanced precision. It is worth noting that these experiments were conducted using an older SELECT COPAS instrument model from 2005. Utilizing a more modern COPAS instrument with an upgraded laminar flow and electronic controls, is expected to further reduce these reported female contamination rates. Taken together, this method allowed us to efficiently and effectively sort a large number of larvae, providing valuable insights into the performance of our sex sorting system.

**Figure 2.**
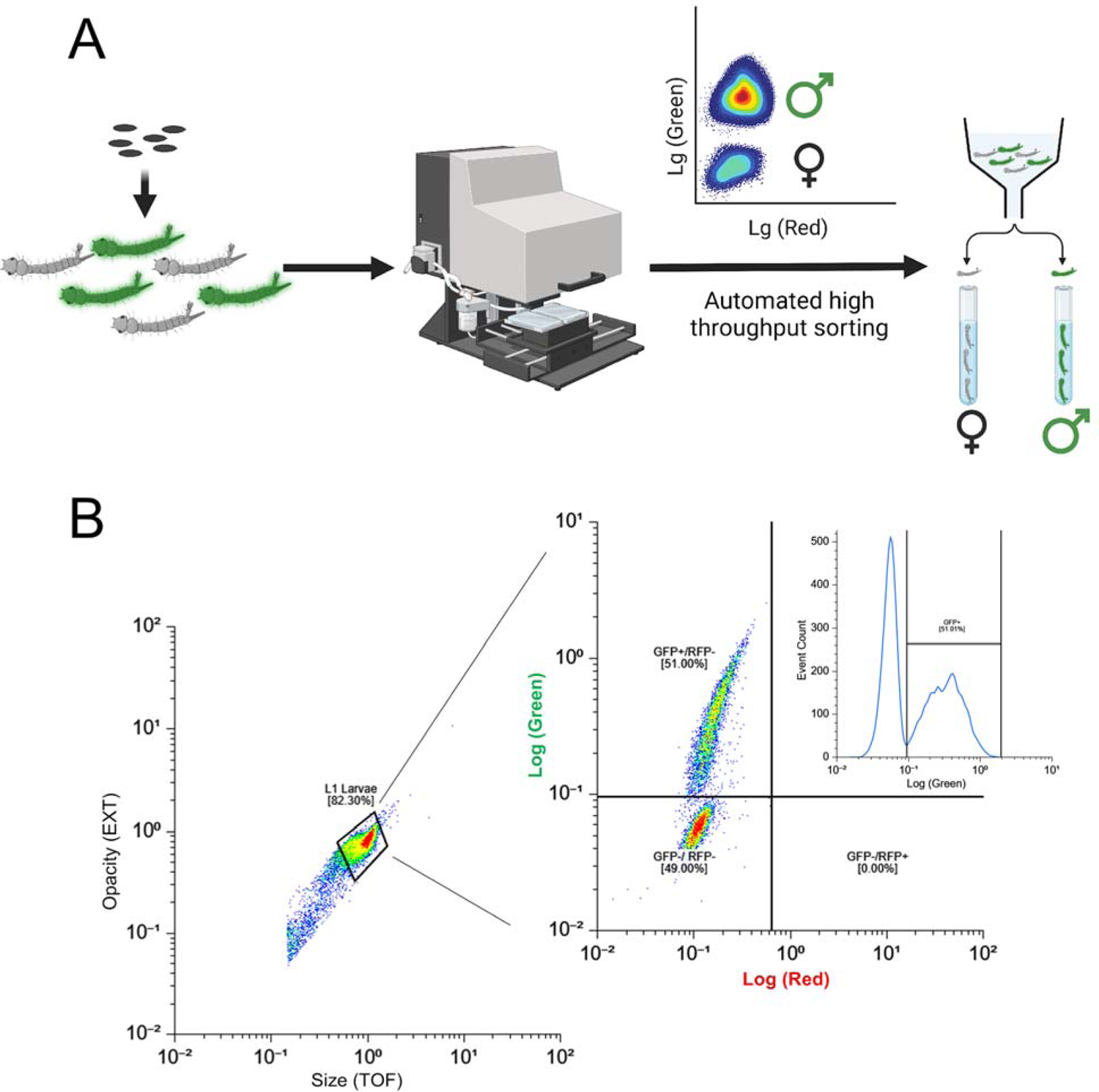
Precise sex-sorting at scale using COPAS. (A) A schematic diagram of large-scale sex-sorting processes using COPAS is shown. The eggs of transgenic mosquitoes, which have been engineered with the SEPARATOR system, were incubated in a vacuum chamber filled with deionized water to hatch them. 24 hours post-egg hatching, the larvae expressing GFP were screened using the COPAS instrument. The GFP-positive larvae were then carefully sorted, and raised in a controlled environment until they reached adulthood. Once the adults have reached maturity, their sexes were verified through various methods. (B) The eggs of transgenic mosquitoes that were genetically engineered to carry the SEPARATOR system were incubated in a vacuum chamber using deionized water. After 24 hours of incubation, the hatched larvae were passed through a COPAS instrument. To ensure accurate sorting, the larvae were selected based on both their opacity and size, and then sorted by the intensity of their GFP expression.

### Sex-enriched genes identification

Previously, sex determination in mosquitoes during the larvae stage has presented challenges, necessitating the reliance on sex-specific morphological characteristics in pupae and adults for precise identification. Consequently, earlier transcriptome analyses primarily concentrated on investigating the sex-related aspects of pupae and adult stages ^55–58^. While there have been some limited studies successfully distinguishing sexes at the L3 and L4 larval stages by assessing the expression of the male determinant factor, Nix, this method requires individual PCR testing of larvae to detect Nix and determine the sex of individual mosquitoes before proceeding with RNA-seq analysis ^57^.

In order to gain valuable insights into the molecular mechanisms controlling sex determination and differentiation during the early developmental stages of *Ae. aegypti* mosquitoes, we employed SEPARATOR to separate male and female L1 larvae. Subsequently, RNA sequencing was conducted to identify genes exhibiting sex-specific expression patterns. Using the data we collected, we conducted a comprehensive analysis of differential gene expression (DGE). Our findings revealed that, at the early L1 larvae stage, 1082 genes exhibited male-enriched expression patterns, while 634 genes exhibited female-enriched expression patterns (**fig. S5C**). Subsequently, the sex-enriched genes were subjected to enrichment analysis. Among the male-enriched genes, two distinct clusters emerged based on Gene Ontology (GO) terms. The first cluster was related to cilium and microtubule formation (including terms such as cilium, axoneme, microtubule-based process, and cell projection), while the second cluster was associated with cuticle formation (**fig. S5D and table S5**). Moreover, among the female-enriched genes, several distinct clusters emerged based on GO terms, including immune response, metabolic processing, and cuticle formation (**fig. S5E and table S5**).

Following RNA-seq analysis of L1 larvae, we compared our transcriptomic data at the L1 larvae stage with previously collected data from the L3, L4, pupae, and adult stages of mosquitoes ^57^. To determine the overlap between the SEPARATOR set and the other six comparisons, we generated an UpSet plot representing shared sex-enriched genes across all developmental stages (**fig. S7-8 and table S9-10**). In our analysis, we have identified a significant number of genes that were not detected in any of the comparisons performed by previous studies ^57^. This finding suggests that some of these genes may represent early expressed genes that are later turned off in subsequent stages, making them difficult to identify in previous studies.

In order to further explore the investigation of sex-enriched genes during various developmental stages of mosquitoes, we performed GO enriched analysis on the list of sex-enriched genes at different developmental stages. However, we observed a limited number of genes displaying sex-enriched patterns specifically in the L3 and L4 larval stages. Furthermore, when analyzing Matthews’s RNA-seq dataset, we found no significant difference in the expression level of a well-known male determinant factor, Nix^58^, between L3 males and L3 females. This discrepancy could potentially be attributed to the lower sequencing depth in Matthews’s RNA-seq data (approximately 7 million reads per sample) compared to our L1 stage dataset (approximately 25 million reads per sample) (**fig. S6 and table S7**). Additionally, to ensure consistency with samples from other stages, we excluded the dataset derived from adult mosquitoes that had been dissected and isolated to obtain independent samples. As a result, we conducted GO enriched analysis on the sex-enriched gene list for the L1 larvae and pupae stages of mosquitoes. Our results indicate a consistent enrichment of genes involved in cilium and microtubule formation in males throughout the developmental stages of mosquitoes, ranging from L1 larvae to late pupae. Notably, cytoskeleton organization-related GO terms were identified during the early to mid pupae stages. Moreover, GO terms associated with spermatid development and sperm DNA condensation were identified during the late pupae stage of mosquito development (**Fig. 3, fig. S9 and table S11**). These results suggest that sperm development is a continuous process from the early developmental stage (L1 larvae) to the late developmental stage (late pupae) in mosquitoes. For female-enriched genes, we observed a notable emphasis on GO terms associated with DNA replication and DNA repair specifically in female pupae (**Fig. 3, fig. S9 and table S11**).

**Figure 3.**
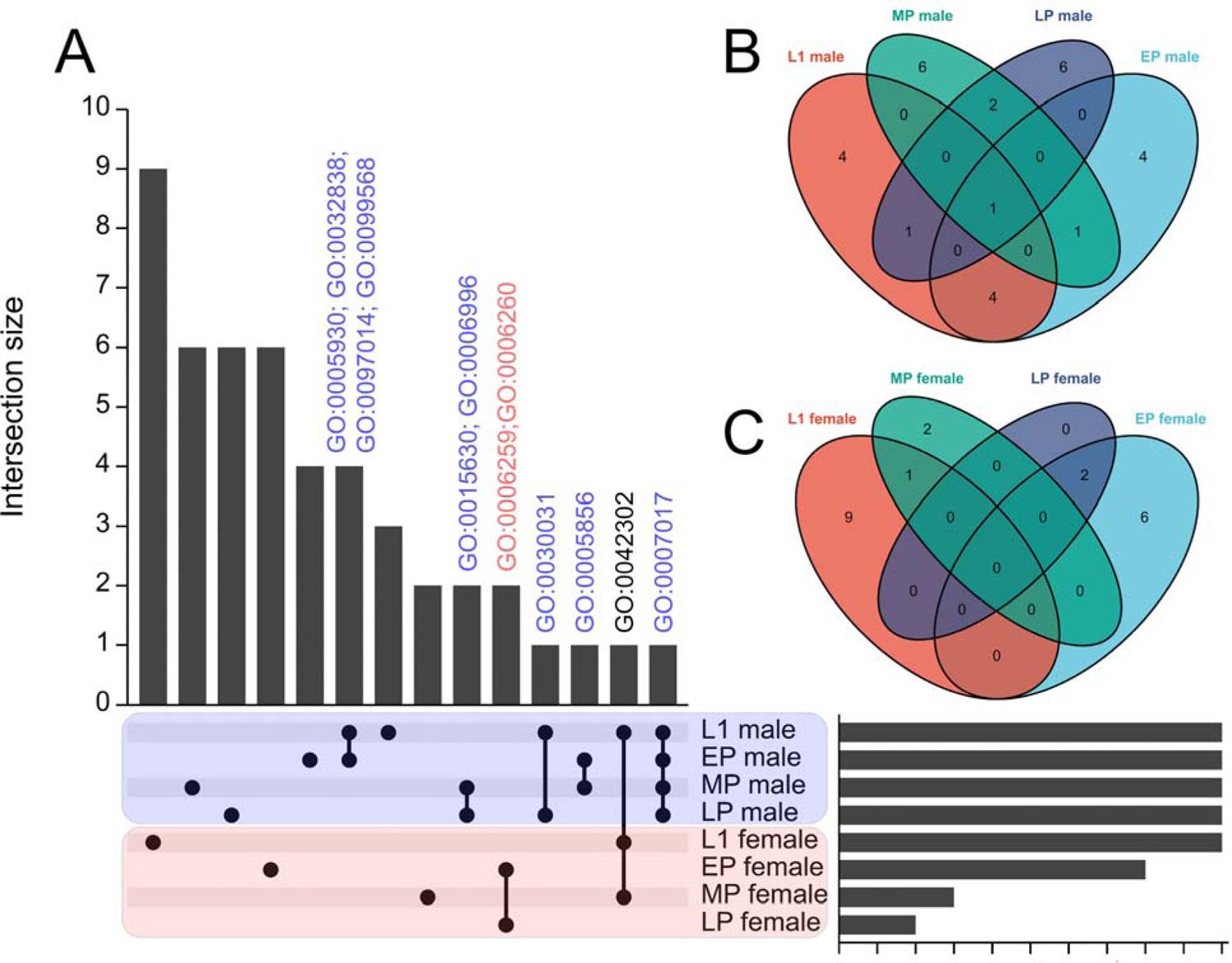
Comparing the transcriptome across larvae and pupae stages of mosquito. SEPARATOR mosquitoes were utilized to individually segregate male and female mosquitoes at the L1 larval stage using the GFP signal. Following separation, total RNA extraction and RNA-seq analysis were performed. The analysis of the early pupae (EP), mid pupae (MP), and late pupae (LP) stages was conducted using data from a previous study. Sexing at the pupae stages relied on sex-specific morphological differences. Sex-enriched genes were identified using DESeq2 and then performed GO enrichment analysis. The shared genes between the L1 stage and all other comparisons were determined, and visualizations such as (A) UpSet plots and (B-C) Venn diagrams were created to represent these shared genes. The common GO terms across various developmental stages of mosquitoes are labeled above the bar. GO:0005930 axoneme; GO:0032838 plasma membrane bounded cell projection cytoplasm; GO:0097014 ciliary plasm; GO:0099568 cytoplasmic region; GO:0015630 microtubule cytoskeleton; GO:0006996 organelle organization; GO:0006259 DNA metabolic process; GO:0006260 DNA replication; GO:0030031 cell projection assembly; GO:0005856 cytoskeleton; GO:0042302 structural constituent of cuticle; GO:0007017 microtubule-based process.

Previously, it was challenging to determine the sex of larvae. As a result, previous transcriptome analyses relied on separating sexes after the pupae stage, which caused the loss of sex-specific samples during early developmental stages. In this study, we utilize our sex-specific RNA-seq results to compare them with a previously collected dataset of variable developmental stage RNA-seq data^56^ (**table S12-13**). To assess the expression of sex-specific genes during early development (prior to the pupae stage), we initially analyzed the genes identified through mfuzz clustering analysis, specifically focusing on those designated as L1 or L2-L4 specific. Cluster 17 consisted predominantly of genes expressed in L1, while cluster 1 encompassed genes expressed in L2-L4 (**fig. S10**). Using a membership cutoff of 0.75, we found that cluster 17 contained 268 genes and cluster 1 contained 383 genes. Among these, 73 (27%) and 134 (35%) were determined to be sex-specifically expressed (**table S14-15**). To expand our investigation beyond the mfuzz clusters, we examined genes that exhibited no expression (TPM values below 1) in carcass, testes, ovary, or pupae, but displayed TPM values above 1 (**table S16**) or 10 (**table S17**) in first instar larvae. These were considered early-expressed genes that were not detected at later stages. We identified 210 and 93 such genes in the respective datasets. Among these, 76 (36%) and 46 (49%) were identified as sex-specifically expressed (**table S16-17**). These findings are highly promising and contribute valuable insights.

In summary, our comparison with the developmental transcriptome data from Akbari et al. demonstrates our ability to identify genes expressed during early developmental stages, such as L1, which was previously unattainable. Furthermore, the comparison with Matthews’ sex-specific data reveals that the differentially expressed genes we have identified exhibit both overlap with known genes and a significant number of new sex-specific candidates, and these may prove useful for future investigation.

## Discussion

Sex separation is crucial in the process of utilizing insects for genetic biocontrol^14^. To make this approach more sustainable and cost-effective, it is important to improve methods for sex separation. In this study, we present a highly efficient method for selecting male larvae in *Aedes* mosquitoes called SEPARATOR. This approach utilizes a COPAS instrument to positively select male L1 larvae that express a dominant male-specific reporter gene (EGFP). This is a positive-selection system, meaning that only those larvae expressing the dominant male-specific GFP are precision sorted. Therefore, if EGFP becomes mutated or breaks, those individuals will not be selected. Moreover, the use of a male spliced intron further increases the robustness since females will not have the machinery to properly splice this intron to generate EGFP.

In comparison to existing techniques, SEPARATOR has several advantages, including its exceptional efficiency, precision, and potential for portability across multiple species. This is due to the evolution conserved sex determination gene, Dsx. In the past, sex sorting relied on morphological differences between male and female mosquitoes during their pupal and adult stages to differentiate between the two sexes. While this method is effective, it does require a significant amount of human effort and time. Unfortunately, this makes it challenging to automate the process on a large scale, which could be a hindrance. Therefore, we need new ways to differentiate between male and female mosquitoes which are faster and more efficient.

In *Aedes*, there is a distinct size difference between female and male pupae, with females being larger. This characteristic allows for pupae to be size-sorted through a sieve, which serves as the first step in sex-sorting. While this method is effective in separating the sexes, up to 2-5% of female pupae can still remain in the male-enriched batch. To further reduce the number of female mosquitoes, a secondary sex-sorting step is employed using an AI-assisted optical system^28, 29^. This system utilizes visual cues from sexual dimorphic differences in adult mosquito morphology to identify and remove any remaining female mosquitoes. This second step significantly reduces female contamination rates to an incredibly low level of 0.02% to 1.131il10^-7^%^59^. However, it should be noted that this step is done at the fragile adult stage, which is not ideal for sorting as adults are very fragile. Furthermore, the COPAS instrument is capable of high-throughput selection, able to sort up to 36K larvae per hour which really makes this a scalable approach.

In *Aedes aegypti* and *albopictus*, transgenic lines ectopically expressing the male determining factor *Nix* have been shown to convert females into males, and enable the positive selection of male mosquitoes ^54, 58, 60^. In *Aedes albopictus*, despite a slight reduction in mating competitiveness, these lines exhibit promise as robust Genetic Sexing Strains (GSS)^54^. Nevertheless, the transformed males were unable to engage in flight in *Ae. aegypti*, mainly due to the absence of *myo-sex*, a closely associated gene found in the M-locus that encodes a flight muscle myosin specific to males^61^. This issue will pose challenges in terms of maintenance and transgene delivery. Interestingly, in *Ae. albopictus*, one or more endogenous copies of *myo-sex*-like genes, which are not linked to the M-locus, are significantly activated in pseudo-males. As a result, pseudo-males of *Ae. albopictus* exhibited proficient flying abilities^54^.

The *transformer-doublesex* splicing cascade has been evolutionarily conserved for over 300 million years, spanning across diverse insect orders like *Diptera*, *Coleoptera*, and *Hymenoptera*. This remarkable preservation strongly suggests that this molecular pathway originated in the ancestral holometabolous insect^62–67^. However, a significant deviation from this pattern is evident in *Lepidoptera*, where the *tra* gene has experienced secondary loss. Consequently, in *Lepidoptera*, the regulation of *dsx* splicing relies on a male-specific protein and a female-specific piRNA^68^. Due to its evolutionary conservation, the *dsx* splicing module (SEPARATOR) can be readily utilized as a genetic sex-sorting strategy across a diverse range of insect species.

There were a few unexpected results associated with SEPARATOR. During the 15 generations reported, no DsRed-expressed mosquitoes were observed. Previous research has shown that in *D. melanogaster* and *Ae. aegypti*, the presence of the fruitless (Fru) protein is not detected in females, despite the expression of the corresponding transcripts ^69–71^. Furthermore, a study conducted in *D. melanogaster* has revealed that the translation of the *fru* female splicing product is suppressed by the Tra protein^70^. This finding suggests that the transcripts specific to females are not translated. In *D. melanogaster*, both Dsx and Fru function as sex-determining factors and are regulated by the Tra protein. Nevertheless, Proving the inability of a gene to undergo translation presents significant challenges. As a result, uncertainty persists regarding whether Dsx follows a comparable mechanism of translation inhibition, emphasizing the necessity for additional investigation. Fortunately, the male-specific product of *AaeDsx* demonstrates successful functionality (**Fig. 1C-E**). Additionally, our objective is to generate a positively selected population of males to enhance reliability and ensure accurate sex sorting. Hence, considering the next generation of SEPARATOR, utilizing it for positive male selection and incorporating a ubiquitous expression system for a transgenic marker could be a viable approach.

SEPARATOR is robust and can be easily adapted for use in both classical Sterile Insect Technique (SIT) and *Wolbachia*-based Insecticide-Treated Insect Technique (IIT) ^21, 25, 26^(**Fig. 4**). The current method for breeding male mosquitoes involves culturing both sexes to the pupal or adult stage for sex sorting, after which the female mosquitoes are removed. In the context of Wolbachia-based IIT, the precise release of only males is crucial. The inadvertent release of females can compromise the effectiveness of the intervention, highlighting the urgent requirement for accurate sexing methods. However, current methods are not as efficient as SEPARATOR. SEPARATOR utilizes less time for male-sorting at the L1 larvae stage, making it a more efficient use of breeding space and feed to raise male mosquitoes. The rapidity of COPAS sorting allows for multiple iterations of screening, effectively minimizing the risk of female contamination. With this system, researchers and breeders can save time and resources by identifying and separating male mosquitoes at an early stage of development. This allows for a more efficient and cost-effective method of breeding and raising male mosquitoes for research and other purposes. Taken together, SEPARATOR provides a valuable tool to help revolutionize the way we sort sexes, making the process more efficient, cost-effective, and reliable and may prove to be a valuable addition to genetic biocontrol programs.

**Figure 4.**
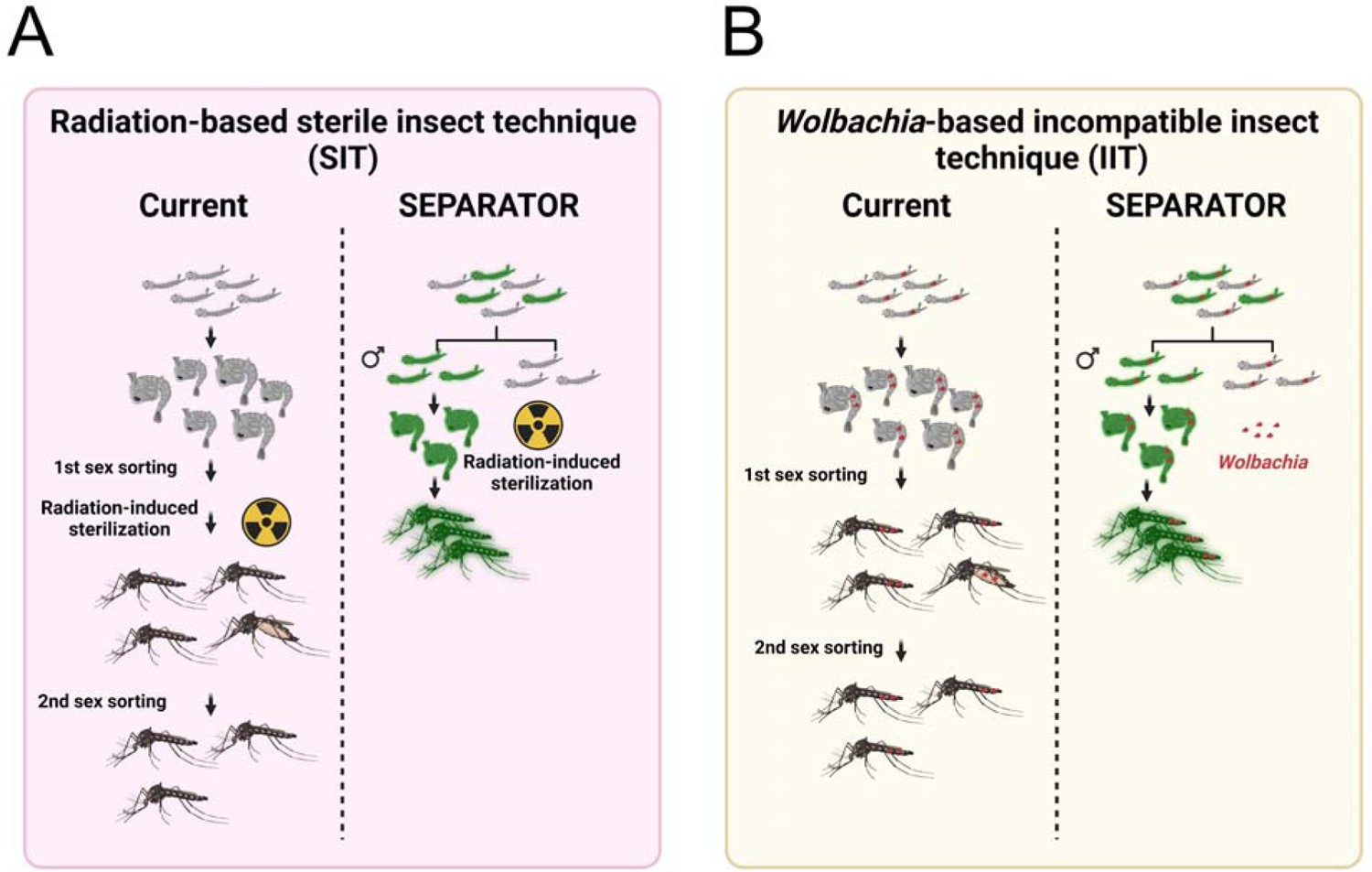
SEPARATOR provides a more simplified and economical sex-sorting of *Ae. aegypti* mosquitoes than the currently available two-step sex-sorting approaches. (A) The process of generating radiation-induced sterile male mosquitoes begins with the cultivation of mosquitoes. The larger female pupae are removed using a sieve. However, it should be noted that some smaller female pupae may still remain after the first sex sorting. Once the mosquitoes have been irradiated, emerging adult mosquitoes are screened out by an image recognition AI that has been trained to discriminate between female and male adult morphologies during the second sex sorting. The SEPARATOR approach utilizes a male-specific reporter (GFP) to positively select male L1 larvae. This can be done using the COPAS instrument, which is capable of high-throughput selection at a speed of up to 10 larvae per second. By removing female larvae early in the development process, SEPARATOR supports a more efficient production of males for SIT application. Furthermore, SEPARATOR allows for the transportation and release of irradiated sex-sorted pupae. This means that adult male mosquitoes can emerge directly into the environment without incurring additional fitness costs from handling and transportation. (B) The *Wolbachia*-based incompatible insect technique (IIT) utilizes a two-step sex-sorting approach for sorting the *Wolbachia*-infected male mosquitoes currently in use. The process of generating *Wolbachia*-infected SEPARATOR mosquitoes is relatively simple, as it involves crossing *Wolbachia*-infected female mosquitoes with SEPARATOR male mosquitoes. The resulting Wolbachia-infected male L1 larvae that express a male-specific reporter (GFP) are then positively selected using the COPAS instrument.

## Materials and Methods

### Molecular Cloning and Transgenesis

To create the endogenous AaeDsx splicing module construct, we amplified the fragment of endogenous exons and introns from the genomic DNA of *Ae. aegypti* using PCR. We then linearized the previous mCherry and EGFP containing plasmid, 1122I, using the restriction enzyme PacI. The linearized 1122I plasmid and the fragment of endogenous exons and introns were used in a Gibson enzymatic assembly method to build the 1174CX plasmid. To open the reading frame of the female-specific product and allow for in-frame expression of the mCherry coding sequence, we substituted the endogenous exon 5b with an engineered exon 5b that had the stop codons removed. The sequence of the engineered exon 5b was synthesized using the gBlocks® Gene Fragment service. We removed the endogenous exon 5b by cutting it with the restriction enzymes PmlI and SnaBI, and then used Gibson assembly to incorporate the engineered exon 5b containing fragment into the cut 1174CX plasmid, resulting in the 1174C plasmid. To generate a plasmid for sex-specific expression of DsRed and EGFP, we amplified the fragment of the engineered exon 5b and DsRed coding sequences from the 1174C plasmid and the previous 874Y plasmid. We then fused these two fragments together using overlapping PCR. Finally, we used omega PCR to substitute the sequence from the mCherry coding sequence to the 3xP3 promoter in the 1174C plasmid with the DsRed coding sequence, resulting in the 1174D plasmid. During each cloning step, we selected single colonies and cultured them in LB medium with ampicillin. We then extracted the plasmids (using the Zymo Research Zyppy plasmid miniprep kit) and performed Sanger sequencing. The final plasmids were maxi-prepped (using the Zymo Research ZymoPURE II Plasmid Maxiprep kit) and fully sequenced by Primordium. All primers are listed in **table S18**. The complete annotated plasmid sequences and plasmid DNA are available at Addgene (ID: 200012). Transgenic lines were created by microinjecting preblastoderm stage embryos with a mixture of the piggyBac plasmid and a transposase helper plasmid. Four days after microinjection, the G0 embryos were hatched and the surviving pupae were separated and sexed. The pupae were placed in separate cages for males and females, along with wild-type male pupae in the female cages and wild-type female pupae in the male cages, in a 5:1 ratio. After several days to allow for development and mating, a blood meal was provided and eggs were collected, aged, and hatched. The larvae with positive fluorescent markers were isolated using a fluorescent stereomicroscope. To isolate separate insertion events, male transformants with fluorescent markers were crossed with female transformants without fluorescent markers, and separate lines were established. The individual genetic sexing lines (1174D) were maintained as mixtures of homozygotes and heterozygotes, with periodic elimination of wild-type individuals. The genetic sexing line (1174D) was homozygosed through approximately ten generations of single-pair sibling matings, selecting individuals with the brightest marker expression each generation.

### Mosquito Rearing and Maintenance

*Ae. aegypti* mosquitoes were obtained from the Liverpool strain, which was previously used to generate the reference genome^57^. These mosquitoes were raised in incubators at 30°C with 20-40% humidity and a 12-hour light/dark cycle in cages (Bugdorm, 24.5 x 24.5 x 24.5 cm). Adults were given 10% (m/V) aqueous sucrose ad libitum, and females were given a blood meal by feeding on anesthetized mice for approximately 15 minutes. Oviposition substrates were provided about 3 days after the blood meal. Eggs were collected, aged for about 4 days to allow for embryonic development, and then hatched in deionized water in a vacuum chamber. Approximately 400 larvae were reared in plastic containers (Sterilite, 34.6 x 21 x 12.4 cm, USA) with about 3 liters of deionized water and fed fish food (TetraMin Tropical Flakes, Tetra Werke, Melle, Germany). For genetic crosses, female virginity was ensured by separating and sexing the pupae under the microscope based on sex-specific morphological differences in the genital lobe shape (at the end of the pupal abdominal segments just below the paddles) before releasing them to eclose in cages. These general rearing procedures were followed unless otherwise noted. In order to increase the number of homozygotes in the 1174D transgenic line, we transferred both the high-intensity GFP pupae and female GFP-negative pupae to a cage and allowed them to mate after eclosion. Female mosquitoes were fed a blood meal, and five adult females were individually transferred to egg tubes for colonization and egg collection. The eggs from each colony were hatched and reared. The colonies with a higher proportion of female EGFP negatives and male EGFP positives were selected for colonization in the next generation. After a few rounds of colonization, the colonies with 100% of males in the strong EGFP positive group and 100% of females in the GFP negative group were selected and propagated for expansion.

### COPAS Fluorescent Sorting, Sexing and Imaging

To determine the precise number of larvae in COPAS clusters, the COPAS raw data was filtered based on the optical density and size measurements of the individuals, “log(EXT)” and “log(TOF),” to remove outliers such as egg debris and dust. Then, a filter was applied based on the individuals’ fluorescence measurements, “log(first fluorescence)” and “log(second fluorescence)”, to further refine the data. Finally, the fluorescence measurements were automatically clustered and denoised using Density-Based Spatial Clustering of Applications with Noise (DBSCAN)^72, 73^. The analysis of COPAS data was conducted using the web-based app, Floreada.io (https://floreada.io/). COPAS sorting was performed largely as described for *Anopheles* larvae^44^. *Aedes* eggs stuck to their egg laying paper were briefly rinsed to eliminate dust and debris, immersed in deionized water in a small container, and their hatching was stimulated under partial vacuum (25% of atmospheric pressure) in a vacuum chamber for 30-60 minutes. They were then incubated overnight at 28°C to maximize larval hatching. On the next day, resulting unfed neonate larvae were transferred to the reservoir of a large particle flow cytometry COPAS SELECT instrument (Union Biometrica, Holliston, MA, USA) equipped with a multiline argon laser (488, 514 nm) and a diode laser (670 nm). Larvae were analyzed and sorted with the Biosort5281 software using a 488 nm filter and the following acquisition parameters: Green PMT 500, Red PMT 600, Delay 8; Width 6, pure mode with superdrops. Flow rate was kept between 20 and 70 objects per second through adjusting of the concentration of larvae in the sample. Larvae identified as males (GFP positive) were dispensed in a Petri dish. In these conditions, sorting speed ranged from 4000 to 7400 larvae in 10 minutes (+ 6 minutes for system initialization and 6 minutes for system cleaning and shutdown), the total number of sorted larvae being limited by the number of available larvae. Sorted larval counts provided by the COPAS software were recorded on sorting. For quality control of sorted larvae, the reservoir and fluidics of the instrument were carefully rinsed and the sorted larvae analyzed by passing them once more in the machine. In some experiments, objects falling outside the GFP positive gate were collected in “Enrich” mode to remove GFP negative contaminants from the pool of GFP positive larvae, and verified by microscopy. Mosquitoes were examined, scored, and imaged using the Leica M165FC fluorescent stereomicroscope equipped with the Leica DMC2900 camera. For higher-resolution images, we used a Leica DM4B upright microscope equipped with a VIEW4K camera. To distinguish between male and female pupae in mosquitoes, we used a microscope to observe the sex-specific morphological differences in the genital lobe shape located at the end of the pupal abdominal segments just below the paddles. This allowed us to ensure that we were selecting both male and female pupae for our experiments.

### Determination of Genome Integration Sites

To determine the transgene insertion sites, we performed Oxford Nanopore genome DNA sequencing. We extracted genomic DNA using the Blood & Cell Culture DNA Midi Kit (Qiagen, Cat. No. / ID: 13343) from 5 adult males and 5 adult females of SEPARATOR, following the manufacturer’s protocol. The sequencing library was prepared using the Oxford Nanopore SQK-LSK110 genomic library kit and sequenced on a single MinION flowcell (R9.4.1) for 72 hrs. Basecalling was performed with ONT Guppy base calling software version 6.4.6 using dna_r9.4.1_450bps_sup model generating 3.03 million reads above the quality threshold of Q≧10 with N50 of 7941 bp and total yield of 11.08 Gb. To identify transgene insertion sites, nanopore reads were mapped to plasmids carrying SEPARATOR (1174D, Addgene as plasmid #200012) using minimap2^74^ and further aligned them to the AaegL5.0 genome (GCF_002204515.2). Subsequently, we calculated the average depth of coverage for the three autosomes and the transgene using samtools and visualized the results in R. The coverage depths for chr1, chr2, and chr3 were determined to be 6.31, 6.30, and 6.08, respectively. Interestingly, the coverage depth for SEPARATOR transgene was notably higher at 16.14. Based on the coverage analysis, it appears that the SEPARATOR transgene is present in three copies (**fig. S3**). By examining the read alignments using Interactive Genomics Viewer (IGV), we were able to determine the exact insertion sites. Notably, we identified three copies of the SEPARATOR constructs sequence. The three integration sites are NC_035109.1:92046983, NC_035108.1:444508475 and NC_035107.1:299022928. This finding aligns with the results of the depth of coverage analysis, further supporting the presence of three insertion sites. The second integration site on NC_035108.1 overlaps with the AAEL005024 gene, which is currently classified as an uncharacterized protein. However, the other two integration sites do not overlap with any known genes. The nanopore sequencing data has been deposited to the NCBI SRA (PRJNA985064).

### RNA sequencing (RNA-seq) analysis

To quantify target gene reduction and expression from transgenes as well as to assess global expression patterns, we performed Illumina RNA sequencing. We extracted total RNA using miRNeasy Tissue/Cells Advanced Mini Kit (Qiagen, Cat. No. / ID: 217604) from 50 GFP-positive male mosquitoes and 50 GFP-negative female mosquitoes at L1 larva stage in biological triplicate (6 samples total), following the manufacturer’s protocol. Genomic DNA was depleted using the gDNA eliminator column provided by the kit. RNA integrity was assessed using the RNA 6000 Pico Kit for Bioanalyzer (Agilent Technologies, Cat. No. / ID: #5067-1513), and mRNA was isolated from ∼1 μg of total RNA using NEBNext Poly(A) mRNA Magnetic Isolation Module (NEB, Cat. No. / ID: E7490). RNA-seq libraries were constructed using the NEBNext Ultra II RNA Library Prep Kit for Illumina (NEB, Cat. No. / ID: E7770) following the manufacturer’s protocols. Briefly, mRNA was fragmented to an average size of 200 nt by incubating at 94°C for 15 min in the first strand buffer. cDNA was then synthesized using random primers and ProtoScript II Reverse Transcriptase followed by second strand synthesis using NEB Second Strand Synthesis Enzyme Mix. Resulting DNA fragments were end-repaired, dA tailed, and ligated to NEBNext hairpin adaptors (NEB, Cat. No. / ID: E7335). Following ligation, adaptors were converted to the “Y” shape by treating with USER enzyme, and DNA fragments were size selected using Agencourt AMPure XP beads (Beckman Coulter #A63880) to generate fragment sizes between 250-350 bp. Adaptor-ligated DNA was PCR amplified followed by AMPure XP bead clean up. Libraries were quantified using a Qubit dsDNA HS Kit (ThermoFisher Scientific, Cat. No. / ID: Q32854), and the size distribution was confirmed using a High Sensitivity DNA Kit for Bioanalyzer (Agilent Technologies, Cat. No. / ID: 5067-4626). Libraries were sequenced on an Illumina NextSeq2000 in paired end mode with the read length of 50 nt and sequencing depth of 25 million reads per library. Base calls and FASTQ generation were performed with DRAGEN 3.8.4. The reads were mapped to the AaegL5.0 genome (GCF_002204515.2) supplemented with SEPARATOR transgene sequences using STAR. On average, ∼97.4% of the reads were mapped. The analysis of RNA-Seq data was performed using an integrated web application called iDEP^75^. TPM values were calculated from counts produced by feature counts and combined (**table S4**). Hierarchical clustering of the data shows that for each genotype, all replicates cluster together, as expected (**fig. S5A and B**). DESeq2 was then used to perform differential expression analyses between male (GFP-positive) and female (GFP-negative) at L1 larvae stage (**fig. S5C**). For each DESeq2 comparison, gene ontology enrichments were performed on significantly differentially expressed genes. (**fig. S5D and E and table S5**). Illumina RNA sequencing data has been deposited to the NCBI-SRA (PRJNA985064). For transcriptome comparing analysis, we acquired 47 files consisting of six developmental stages (L3 larvae, L4 larvae, early pupae, mid pupae, late pupae, and adult carcass) from SRA (**table S6**). These files were then aligned to the AaegL5 genome (GCF_002204515.2) using STAR. The analysis of RNA-Seq data was performed using an integrated web application called iDEP^75^. TPM values were calculated from counts produced by feature counts and combined (**table S8**).

## Statistical analysis

Statistical analysis was performed in Prism9 for macOS by GraphPad Software, LLC. At least three biological replicates were used to generate statistical means for comparisons.

## Data availability

Complete sequence maps and plasmids are deposited at Addgene.org (#200012). All Illumina and Nanopore sequencing data has been deposited to the NCBI-SRA (PRJNA985064). All data used to generate figures are provided in the Supplementary Materials/Tables. Generated transgenic lines are available upon request to O.S.A.

## Acknowledgments

We thank Ming Li, Judy Ishikawa, Sammy Lee, and Dylan Turksoy for helping with mosquito husbandry and microinjections. This work was supported by funding from an NIH awards (R01AI151004, R01GM132825, RO1AI148300, RO1AI175152, DP2AI152071), EPA STAR award (RD84020401), and an Open Philanthropy award (309937-0001) awarded to O.S.A. E.M. received funding from Agence Nationale de la Recherche through grant #ANR-19-CE35-0007 GDaMO. The views, opinions, and/or findings expressed are those of the authors and should not be interpreted as representing the official views or policies of the U.S. government. Figures were created using www.BioRender.com.

## Author Contributions

O.S.A and S.C.W. conceptualized and designed experiments; S.C.W. performed molecular analyses, and genetic experiments; I.A. performed bioinformatics: S.C.W, analyzed and compiled the data. E.M. performed the COPAS experiments and analysis. All authors contributed to writing and approved the final manuscript.

## Ethical conduct of research

All animals were handled in accordance with the Guide for the Care and Use of Laboratory Animals as recommended by the National Institutes of Health and approved by the UCSD Institutional Animal Care and Use Committee (IACUC, Animal Use Protocol #S17187) and UCSD Biological Use Authorization (BUA #R2401).

---

**Fig S1.**
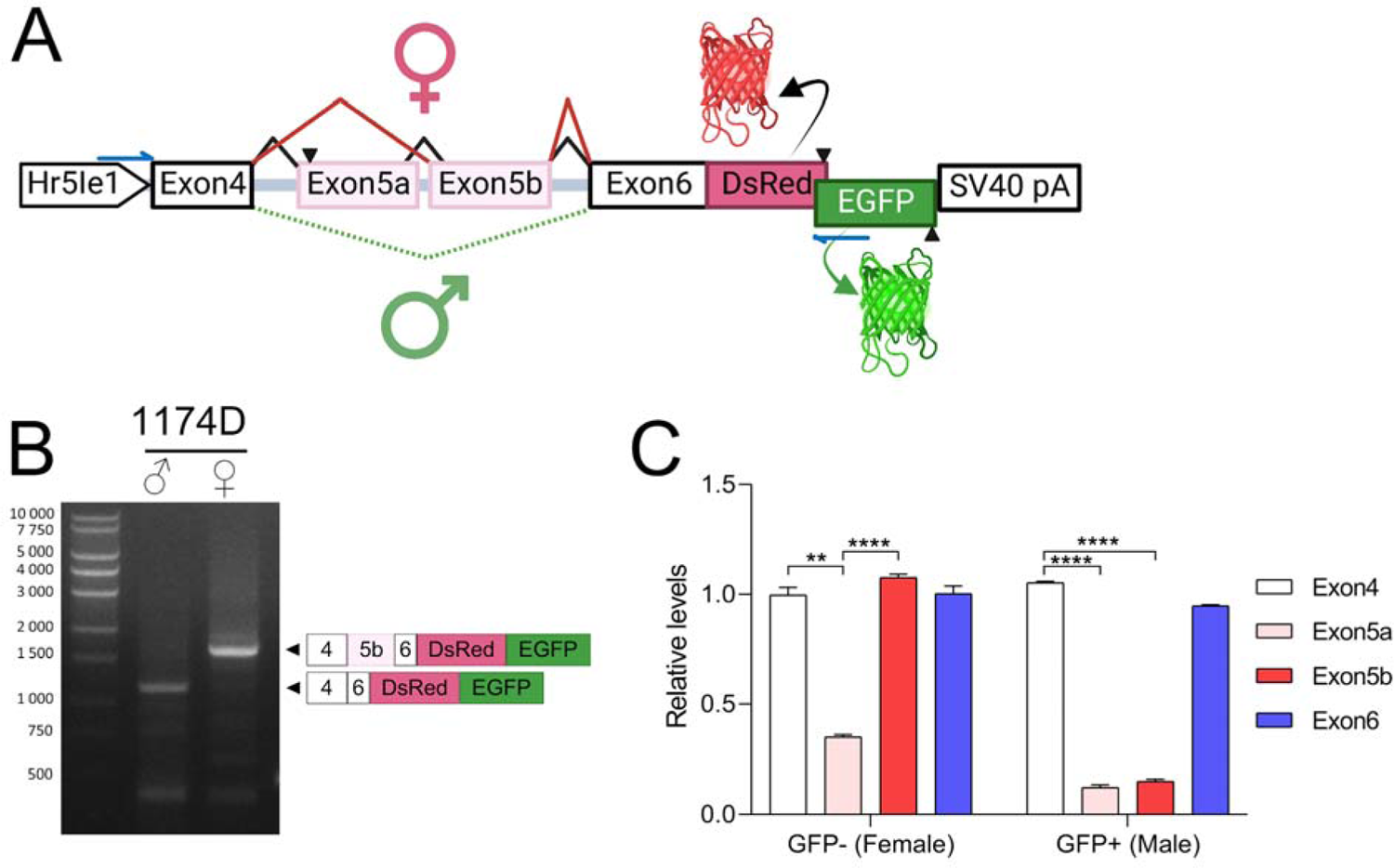
Sex-specific RNA splicing of SEPARATOR. Fifty GFP-positive larvae and fifty GFP-negative larvae at the L1 stage were carefully sorted, and total RNA was extracted from each group. To determine the splicing patterns, RT-PCR was performed using specific primers targeting the 3’ end of the Hr5Ie1 promoter sequence and the 5’ end of the EGFP coding sequence. (A) The relative locations of the primer target sites are indicated by blue arrows. (B) The PCR products were subsequently purified and subjected to sequencing in order to validate the splicing junctions. The resulting splicing patterns are depicted in the right panel. (C) The relative levels of non-sex-specifically regulated exons (exon4 and exon6) and female-specific exons (exon5a, exon5b) of SEPARATOR were determined through RNA sequencing (RNAseq) analysis. The FPKM (fragments per kilobase per million mapped reads) values of each exon were normalized using the average FPKM of the non-sex-specifically regulated exons (exon4 and exon6). The bar plot displays the means and ± SD (standard deviation) for triple biological replicates. Statistical significance of mean differences was assessed using a Tukey’s multiple comparisons test, with p-values denoted as follows: p < 0.01** and p < 0.0001****.

**Fig S2.**
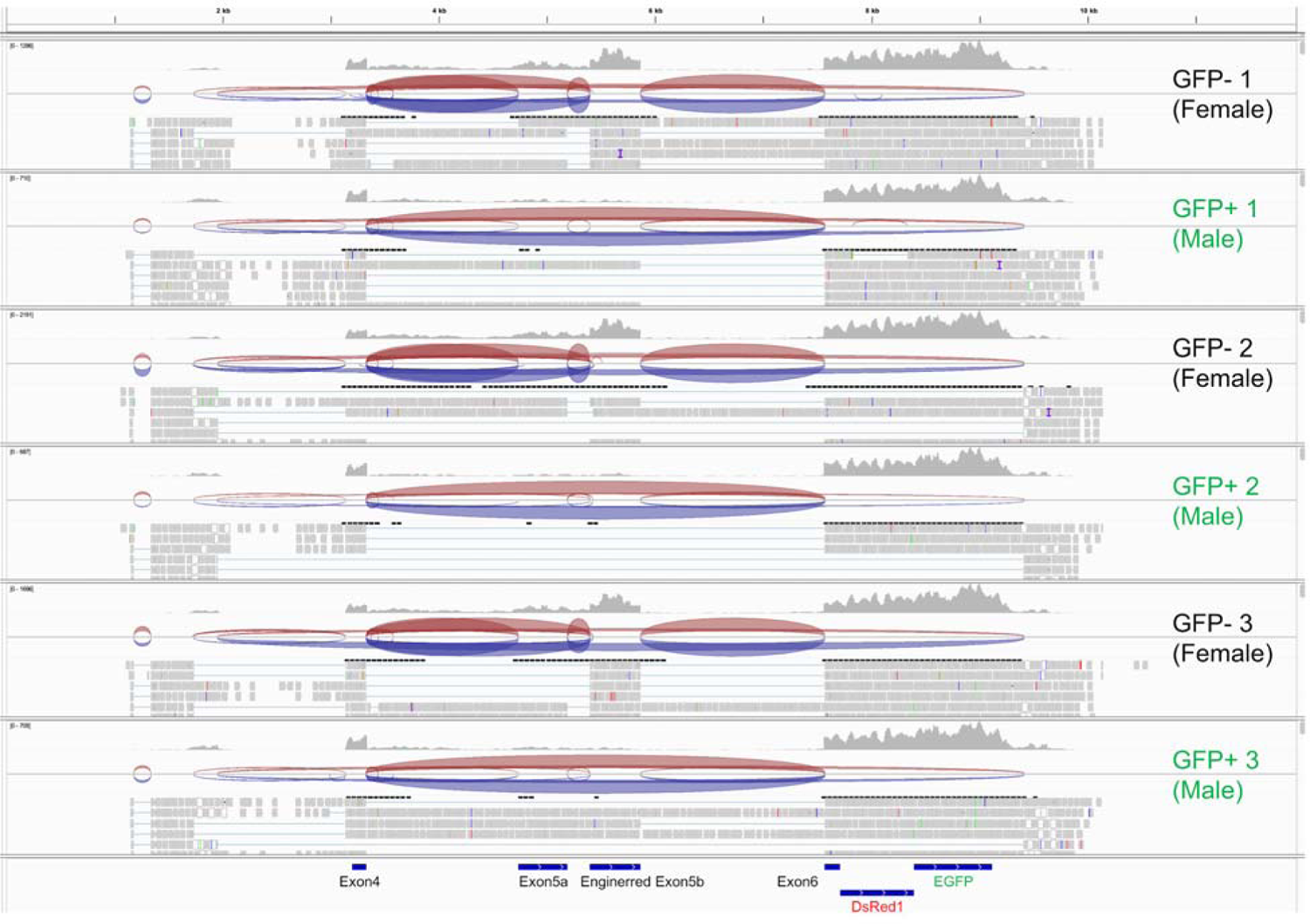
Sex-specific RNA splicing patterns verified through RNA sequencing analysis. The splicing patterns of SEPARATOR were verified through RNAseq analysis in both GFP-positive and GFP-negative mosquitoes, with triple biological replicates for each condition. The RNAseq reads for the different genotypes were aligned, and the location of exons is indicated at the bottom in blue.

**Fig S3.**
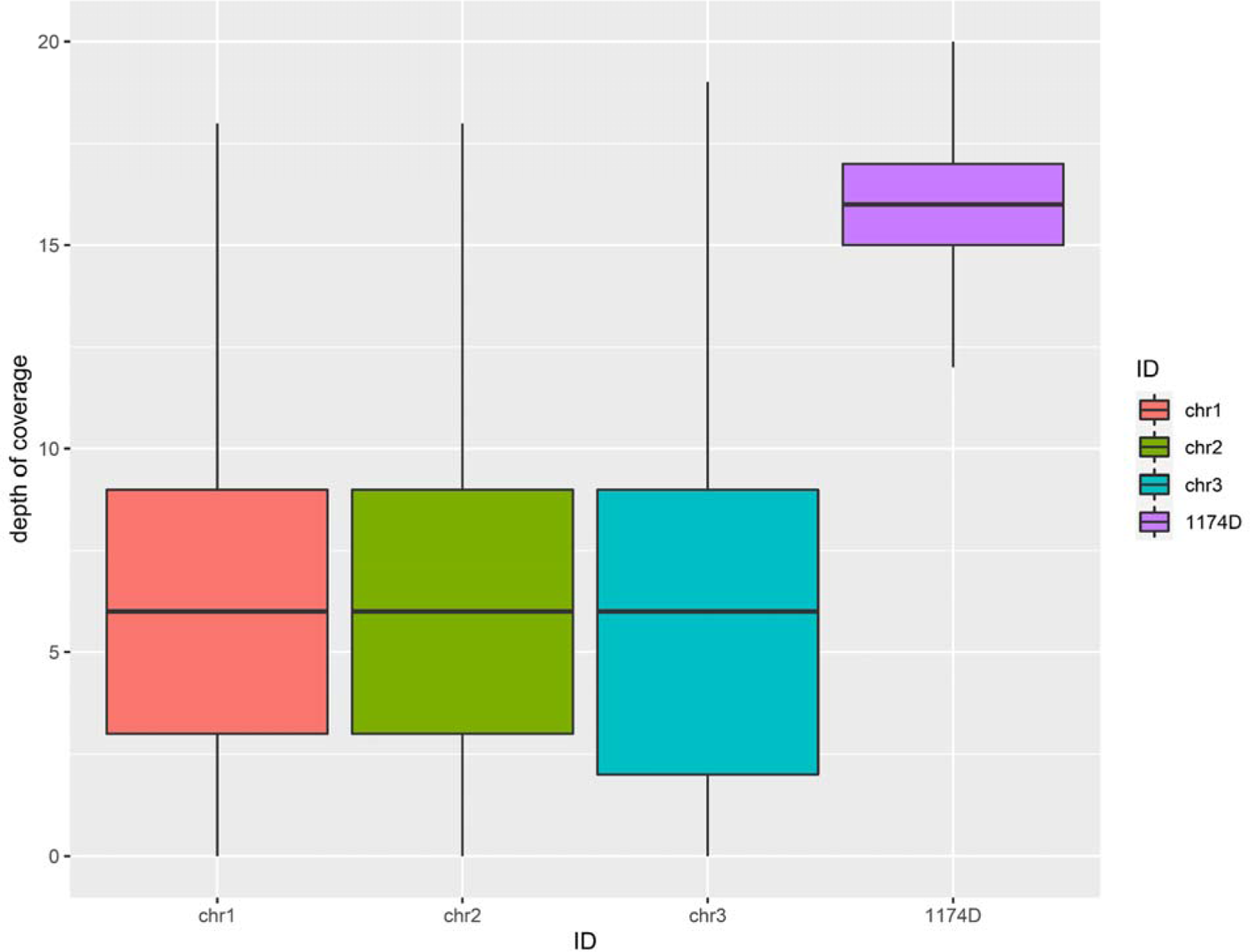
The transgene copy number for SEPARATOR was determined using Oxford Nanopore genome sequencing. A standard box plot is used to illustrate the coverage distributions of three chromosomes (Chr1, Chr2 and Chr3) and the SEPARATOR transgenes (1174D) in SEPARATOR mosquitoes. The center line represents the median, while the first and third quartiles define the boundaries of the box. The upper and lower whiskers extend from the box to the highest and lowest observed values, respectively, but no further than 1.5 times the Interquartile Range (IQR) from the box. Based on the sequencing depths, the coverage for chromosomes 1, 2, and 3 were 6.31, 6.30, and 6.08, respectively, while the coverage for the SEPARATOR transgenes was 16.14. From the coverage analysis, it suggests that the SEPARATOR transgene (1174D) is present in three copies.

**Fig S4.**
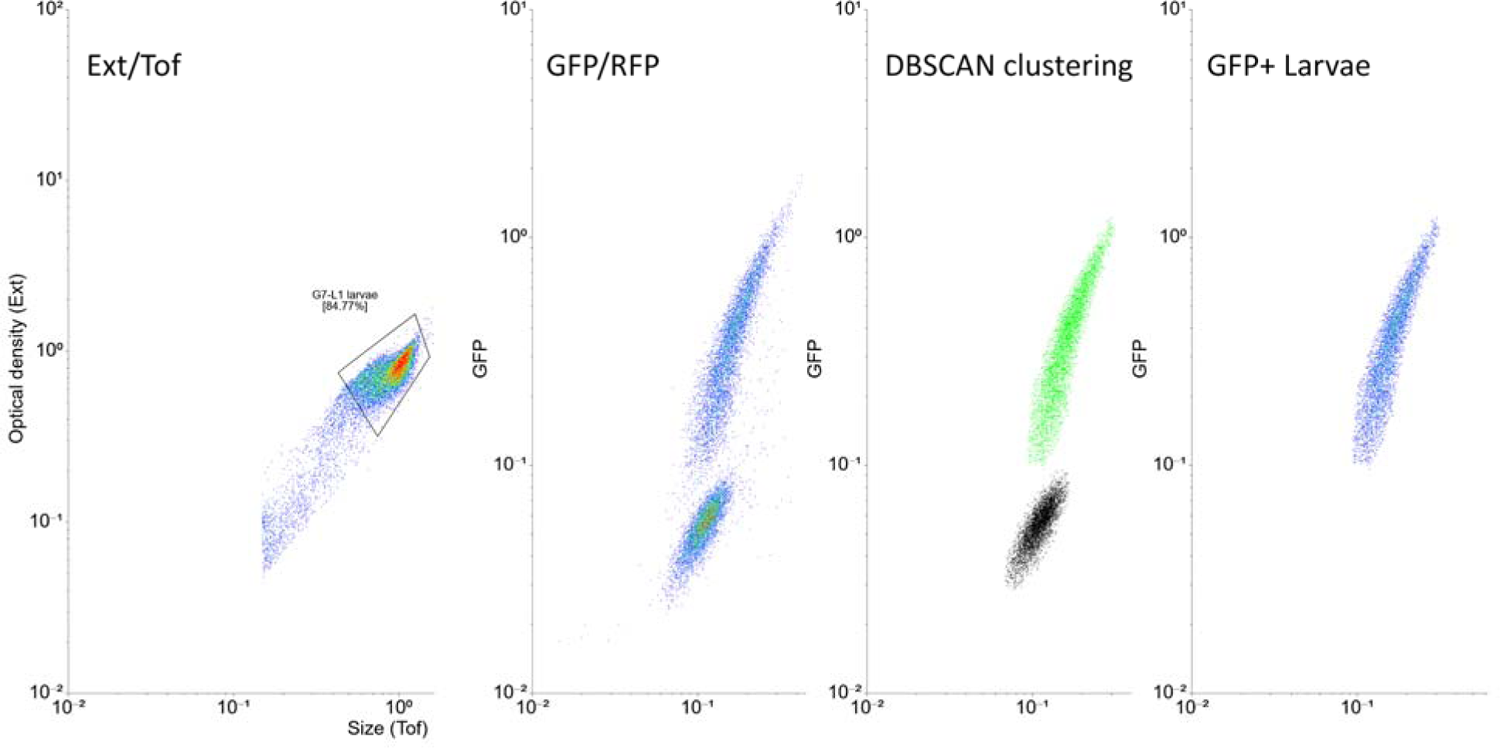
COPAS data processing. The COPAS raw data was initially filtered for the larvae using size and optical density criteria (Ext/Tof). Next, the particles that exhibited fluorescence (GFP/RFP) were gated. Subsequently, DBSCAN clustering was used to automatically cluster and denoise the data. Finally, the larvae that were GFP-positive were selected.

**Fig S5.**
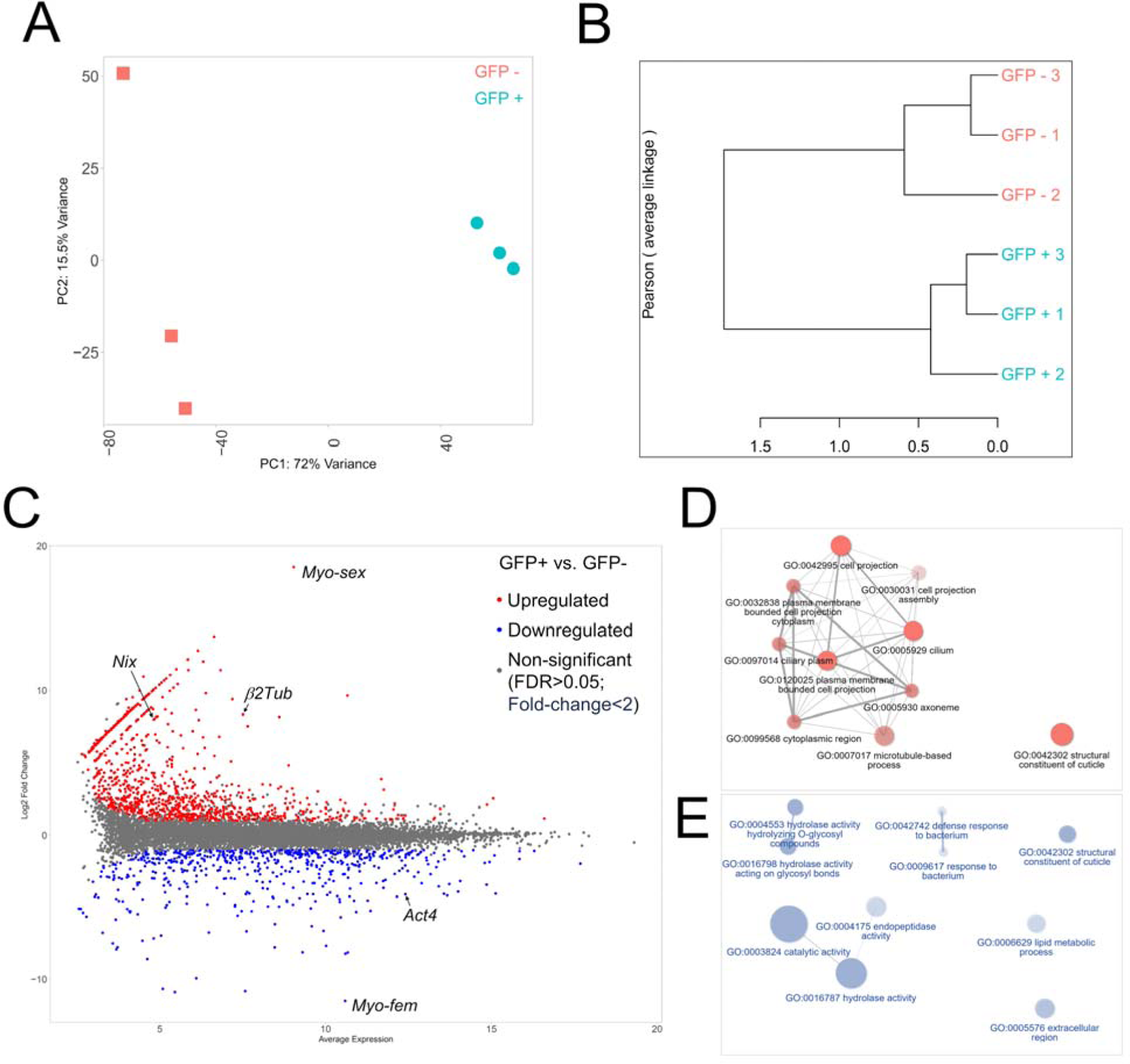
Transcription profiling and expression analysis of GFP-positive and GFP-negative larvae at the L1 stage in SEPARATOR mosquitoes. (A) PCA analysis and (B) hierarchical clustering of six samples used for RNA sequencing. (C) MA-plots were generated to visualize the differential expression patterns between GFP-positive and GFP-negative larvae at the L1 stage in SEPARATOR mosquitoes. In the plot, significantly upregulated (male-enriched) genes (FDR < 0.05 and fold-change > 2) are indicated by red dots, significantly downregulated (female-enriched) genes (FDR < 0.05 and fold-change > 2) are indicated by blue dots, and non-significantly differentially expressed genes are represented by gray dots (FDR > 0.05 or fold-change < 2). Additionally, five well-known sex-enriched genes were marked in the plot. A network visualization was created to illustrate the relationship among enriched Gene Ontology (GO) terms for the upregulated (D) and downregulated (E) genes.

**Fig S6.**
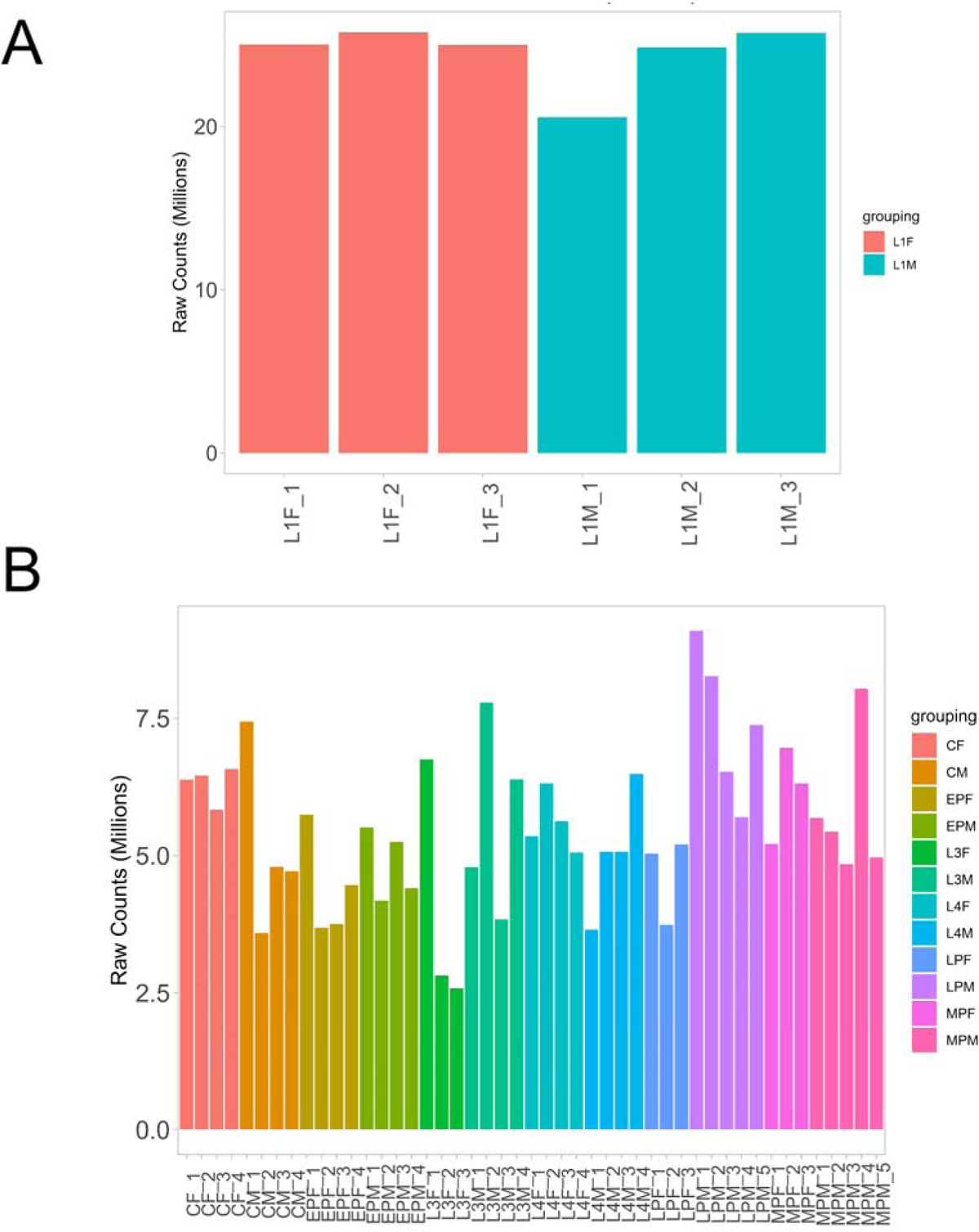
The RNA sequencing depth in SEPARATOR mosquitoes is greater than that observed in previous Matthews’s RNA-seq datasets. In the transcriptome comparison analysis, we employed GFP-positive (Male, L1M) and GFP-negative (Female, L1F) larvae at the L1 stage from SEPARATOR mosquitoes. Additionally, we utilized larvae at the L3 (L3M and L3F) and L4 (L4M and L4F) stages, as well as early pupae (EPM and EPF), mid pupae (MPM and MPF), late pupae (LPM and LPF), and carcass of adult mosquitoes (CM and CF) from Matthews’s RNA-seq datasets. The sequencing depth of the RNA-Seq data was achieved through the utilization of an integrated web application known as iDEP.

**Fig S7.**
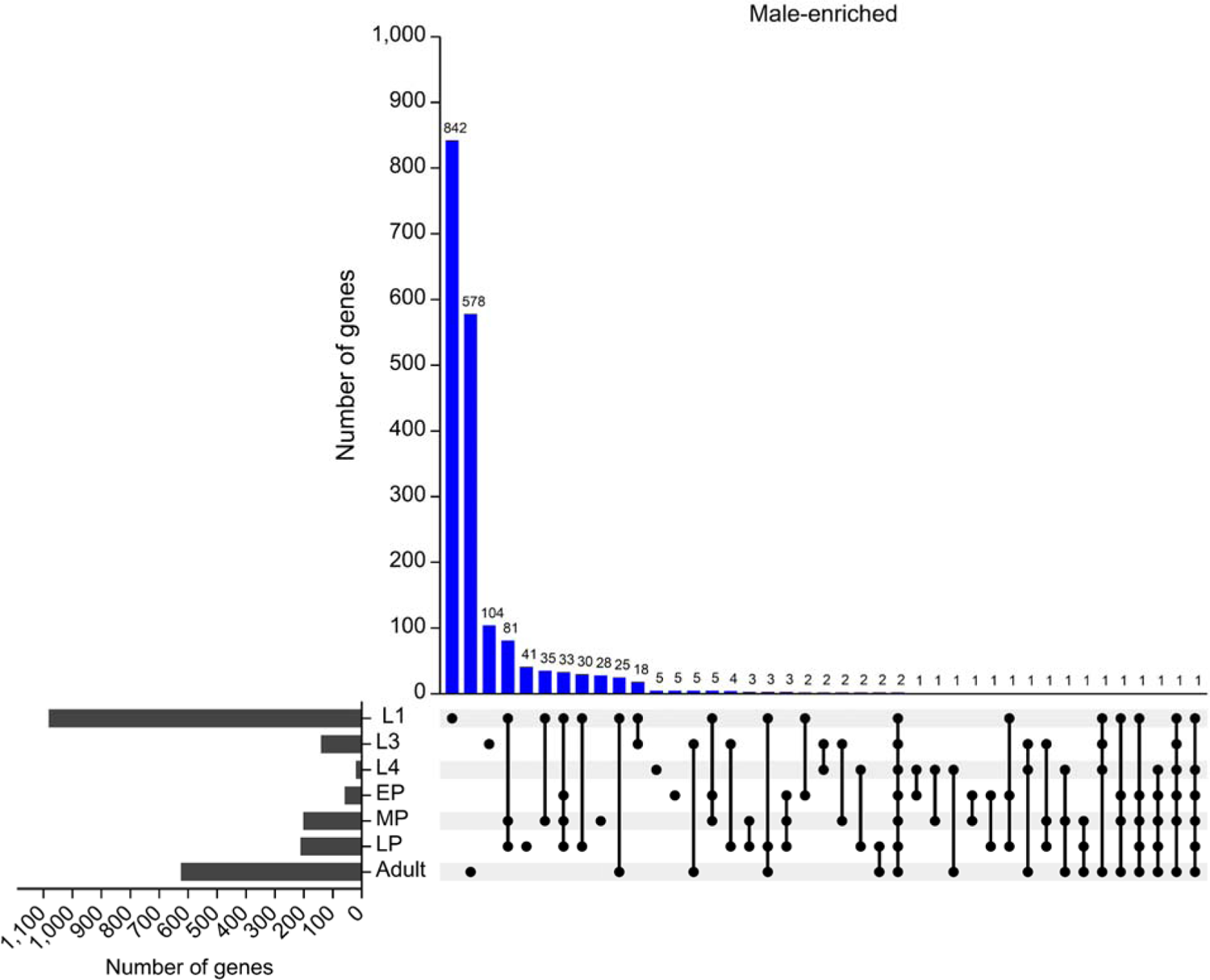
Male-enriched genes from different developmental stages were identified in the transcriptome comparison analysis. In our transcriptome comparison analysis, we included L1 stage larvae from SEPARATOR mosquitoes. Furthermore, we incorporated L3 and L4 stage larvae, along with early pupae (EP), mid pupae (MP), late pupae (LP), and adult mosquito carcass (Adult) from Matthews’s RNA-seq datasets. The correlation of male-enriched genes in this analysis was visualized using an UpSet plot, facilitated by the integrated web application iDEP.

**Fig S8.**
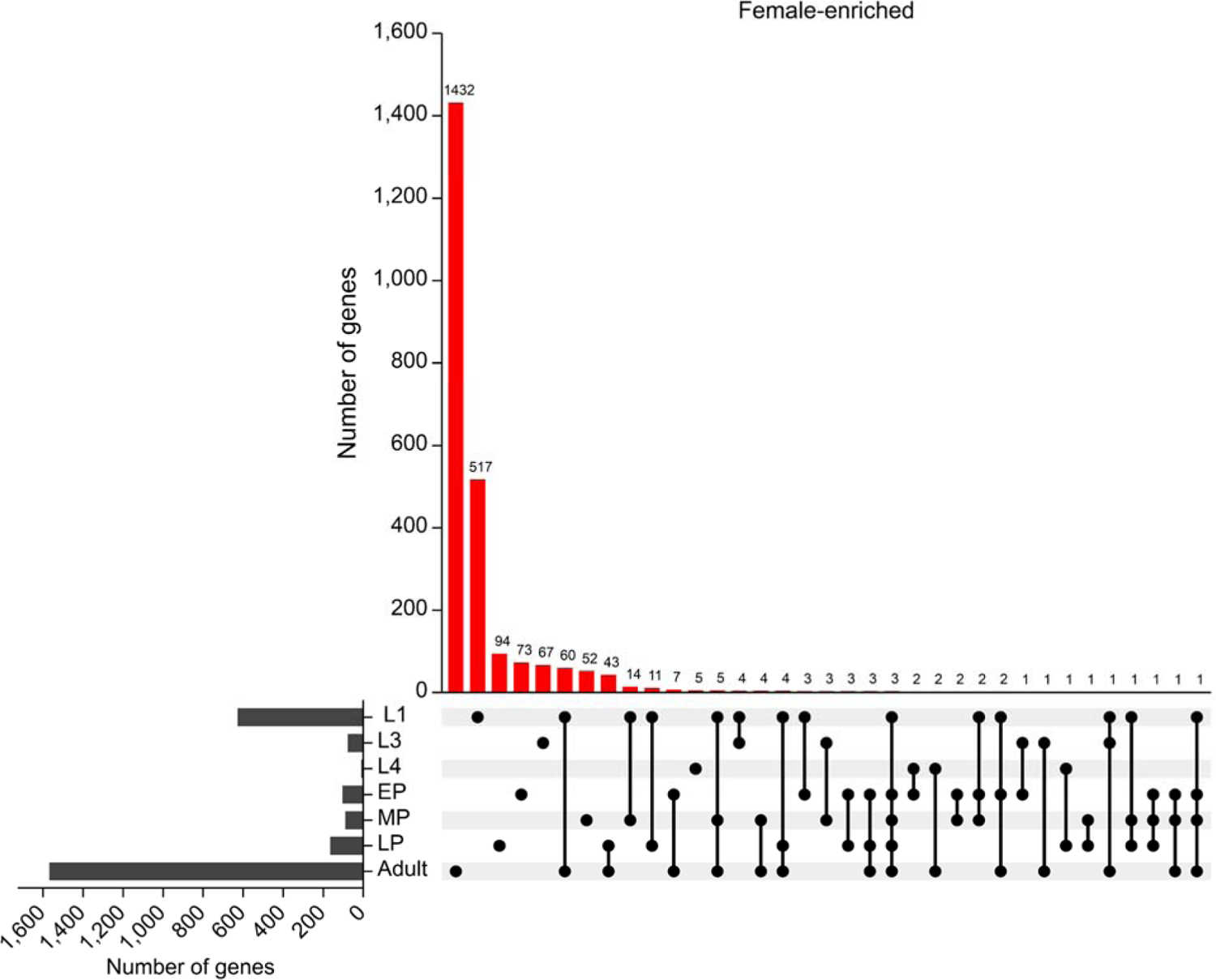
Female-enriched genes from different developmental stages were identified in the transcriptome comparison analysis. In our transcriptome comparison analysis, we included L1 stage larvae from SEPARATOR mosquitoes. Furthermore, we incorporated L3 and L4 stage larvae, along with early pupae (EP), mid pupae (MP), late pupae (LP), and adult mosquito carcass (Adult) from Matthews’s RNA-seq datasets. The correlation of female-enriched genes in this analysis was visualized using an UpSet plot, facilitated by the integrated web application iDEP.

**Fig S9.**
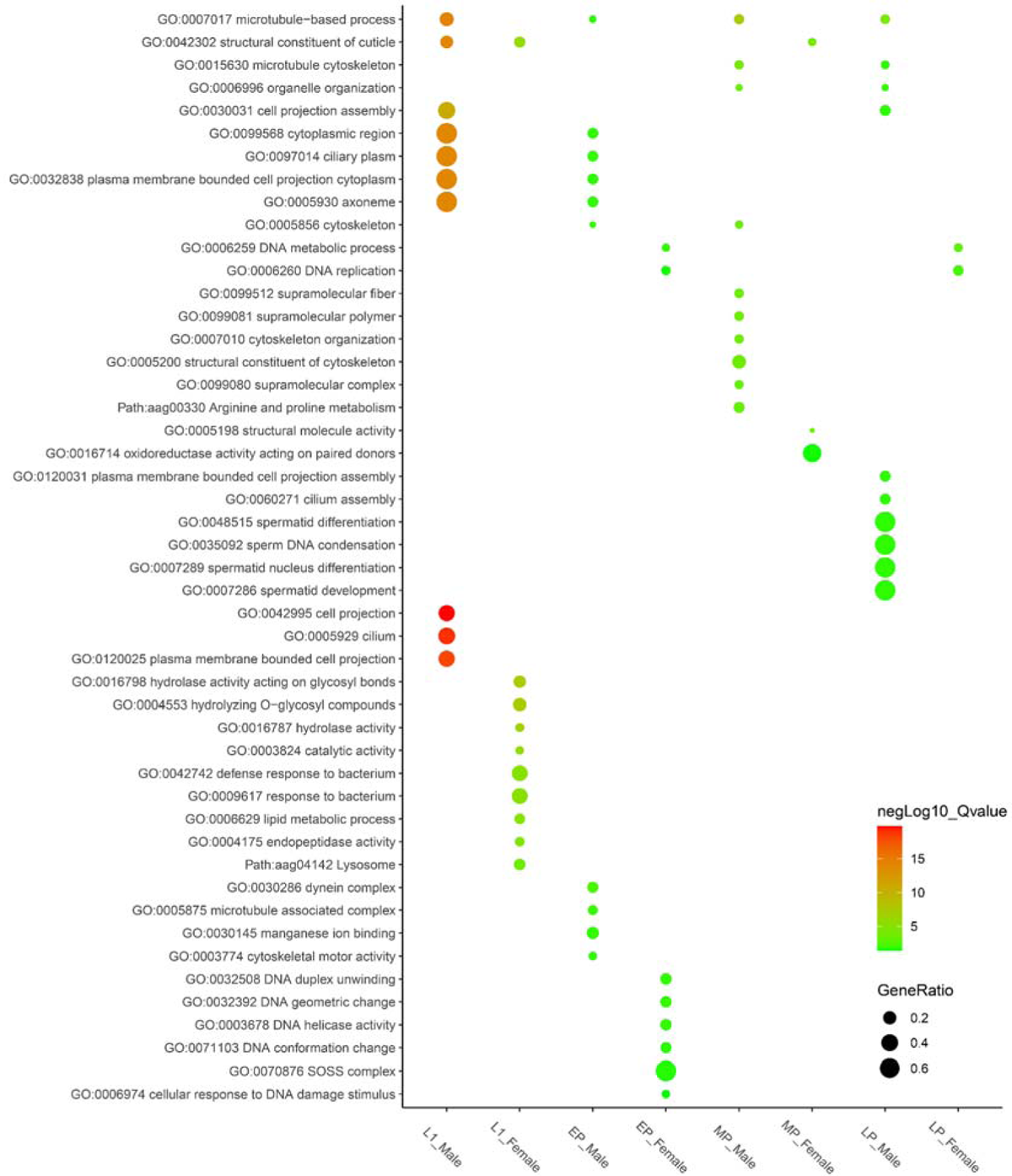
Conducting a gene ontology (GO) analysis on sex-enriched genes throughout various developmental stages In our transcriptome comparison analysis, we included L1 stage larvae from SEPARATOR mosquitoes. Furthermore, we incorporated early pupae (EP), mid pupae (MP), and late pupae (LP), from Matthews’s RNA-seq datasets. The correlation of GO terms in sex-enriched genes within this analysis was identified and facilitated by the integrated web application iDEP.

**Fig S10.**
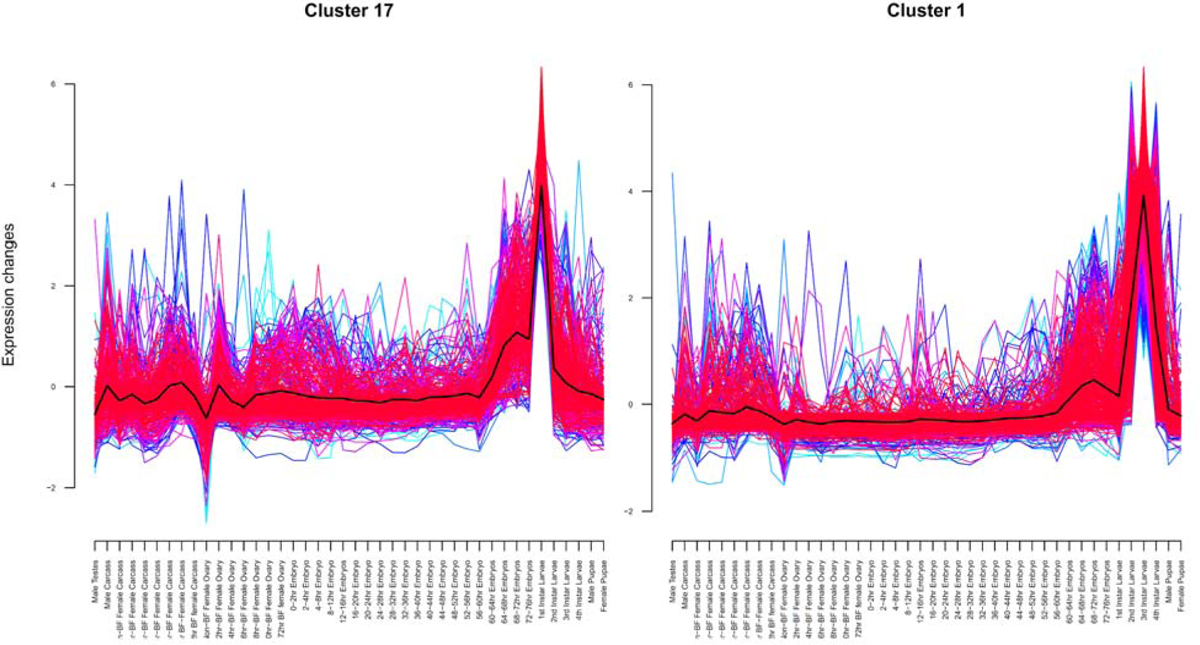
Gene expression analysis and clustering methods are used to identify and isolate a distinct set of genes associated with the larvae stage. Through mfuzz clustering analysis using comprehensive developmental stage data from a previous study, specific genes associated with either L1 or L2-L4 stages were identified. Notably, cluster 17 predominantly consisted of genes expressed in L1, while cluster 1 exhibited gene expression primarily in L2-L4 stages.

## Supplementary Tables

**table S1.** The sex sorting of SEPARATOR mosquitoes during the initial 15 generations

**table S2.** Summary of results from COPAS sorting experiments

**table S3.** FPKM of each exons in SEPARATOR mosquitoes

**table S4.** Combined annotations exons count

**table S5.** Enriched Gene Ontology (GO) terms for GFP-positive versus GFP-negative larvae in SEPARATOR mosquitoes

**table S6.** Library info_matthews

**table S7.** Read count_matthews

**table S8.** Read count_matthews

**table S9.** Upset Result-Male enriched

**table S10.** Upset Result-Female enriched

**table S11.** Upset Result-GO term

**table S12.** Akbari samples pruned

**table S13.** Akbari combined count

**table S14.** deseq2 1174D_pos_vs_neg.annotations.cluster_17.significantly_changed

**table S15.** deseq2 1174D_pos_vs_neg.annotations.cluster_1.significantly_changed

**table S16.** eseq2_1174D_pos_vs_neg.annotations.1st_instar_1tpm.significantly_changed

**table S17.** eseq2_1174D_pos_vs_neg.annotations.1st_instar_10tpm.significantly_changed

**table S18.** Sequences of primers and gBlock fragment used in this study

